# Multiple layers of cryptic genetic variation underlie a yeast complex trait

**DOI:** 10.1101/465278

**Authors:** Jonathan T. Lee, Alessandro L.V. Coradini, Amy Shen, Ian M. Ehrenreich

**Affiliations:** Molecular and Computational Biology Section, Department of Biological Sciences, University of Southern California, Los Angeles, CA 90089-2910, USA; Laboratory of Genomics and Bioenergy, Department of Genetics and Evolution, Institute of Biology, UNICAMP, Campinas, São Paulo, 13083-970, Brazil

**Keywords:** Cryptic genetic variation, epistasis, genotype-environment interaction, complex traits, genetic architecture

## Abstract

Cryptic genetic variation may be an important contributor to heritable traits, but its extent and regulation are not fully understood. Here, we investigate the cryptic genetic variation underlying a *Saccharomyces cerevisiae* colony phenotype that is typically suppressed in a cross of the lab strain BY4716 (BY) and a derivative of the clinical isolate 322134S (3S). To do this, we comprehensively map the trait’s genetic basis in the BYx3S cross in the presence of three different genetic perturbations that enable its expression. This allows us to detect and compare the specific loci harboring cryptic genetic variants that interact with each perturbation. In total, we identify 21 loci, all but one of which interacts with just a subset of the perturbations. Beyond impacting which loci contribute to the trait, the genetic perturbations also influence the extent of additivity, epistasis, and genotype-environment interaction among the detected loci. Additionally, we show that the single locus interacting with all three perturbations corresponds to the coding region of the cell surface gene *FLO11*. Nearly all of the other loci influence *FLO11* transcription in *cis* or *trans*. However, the perturbations reveal cryptic genetic variation in different pathways and sub-pathways upstream of *FLO11*, suggesting that multiple layers of cryptic genetic variation with highly contextual effects underlie the trait. Our work demonstrates an abundance of cryptic genetic variation in transcriptional regulation and illustrates how this cryptic genetic variation complicates efforts to study the relationship between genotype and phenotype.

**Article Summary:** To better understand cryptic genetic variation, we comprehensively map the genetic basis of a trait that is suppressed in a yeast cross. By examining how three different genetic perturbations give rise to the trait, we identify 21 loci harboring cryptic genetic variants, nearly all of which play roles in the transcriptional regulation of the same key gene. We examine these loci in greater detail and show that the perturbations affect every aspect of the trait’s genetic architecture, including which cryptic genetic variants have phenotypic effects, as well as the extent of additivity, epistasis, and genotype-environment interaction among these variants.

## Introduction

Most research on complex traits focuses on characterizing the genetic basis of phenotypic diversity that is visible within populations (Atwell *et al.* 2010; Aylor *et al.* 2011; Mackay *et al.* 2012; Bloom *et al.* 2013). Yet, these same populations can also harbor cryptic genetic variation that does not typically impact phenotype, and is only observable when particular genetic or environmental perturbations occur (Rutherford AND Lindquist 1998; Queitsch *et al.* 2002; Bergman AND Siegal 2003; Dworkin *et al.* 2003; Gibson AND Dworkin 2004; Jarosz AND Lindquist 2010; Geiler-Samerotte *et al.* 2016). This cryptic variation may be an important source of phenotypic variability in medically and evolutionarily significant traits (Le Rouzic and Carlborg 2008; Mcguigan AND Sgro 2009; Paaby AND Rockman 2014). Thus, it is imperative that we determine the extent of cryptic variation within populations, as well as the mechanisms that convert this cryptic variation between silent and visible states. However, such work is inherently difficult because the specific perturbations needed to uncover cryptic variation, as well as the exact identities of the cryptic genetic variants (hereafter, ‘cryptic variants’) that are affected by these perturbations, are rarely known. Such information is critical to obtaining a more complete, mechanistic understanding of cryptic variation.

In previous papers, we developed an experimental system that can be used to systematically identify cryptic variants influencing a colony phenotype in *Saccharomyces cerevisiae* (Taylor AND Ehrenreich 2014; Taylor AND Ehrenreich 2015b; Lee *et al.* 2016; Taylor *et al.* 2016). The lab strain BY4716 (BY), a haploid derivative of the clinical isolate 322134S (3S), and their haploid recombinant progeny form ‘smooth’ colonies when grown on solid media (**Figure 1A**). However, certain *de novo* or induced mutations, as well as recombination between the promoter and coding region of the cell surface gene *FLO11*, can enable some BYx3S segregants to express an alternative ‘rough’ colony phenotype (**Figure 1B**). Throughout the current paper, we refer to these new alleles that are not naturally present in BY or 3S, but make it possible for the rough phenotype to be expressed in the BYx3S cross as genetic perturbations (or ‘GPs’). These GPs on their own are insufficient to cause expression of the rough phenotype; rather, cryptic variants that segregate between BY and 3S are also needed. Moreover, BYx3S segregants that express the trait in the presence of a given GP usually exhibit the phenotype in a temperature-sensitive manner. However, by backcrossing these segregants and examining their backcross progeny across temperatures, additional cryptic variation can often be found that reduces or eliminates temperature sensitivity (**Figure 1C**). In other words, expression of the trait results from complex interactions between the GPs, segregating loci, and temperature.

**Figure 1:**
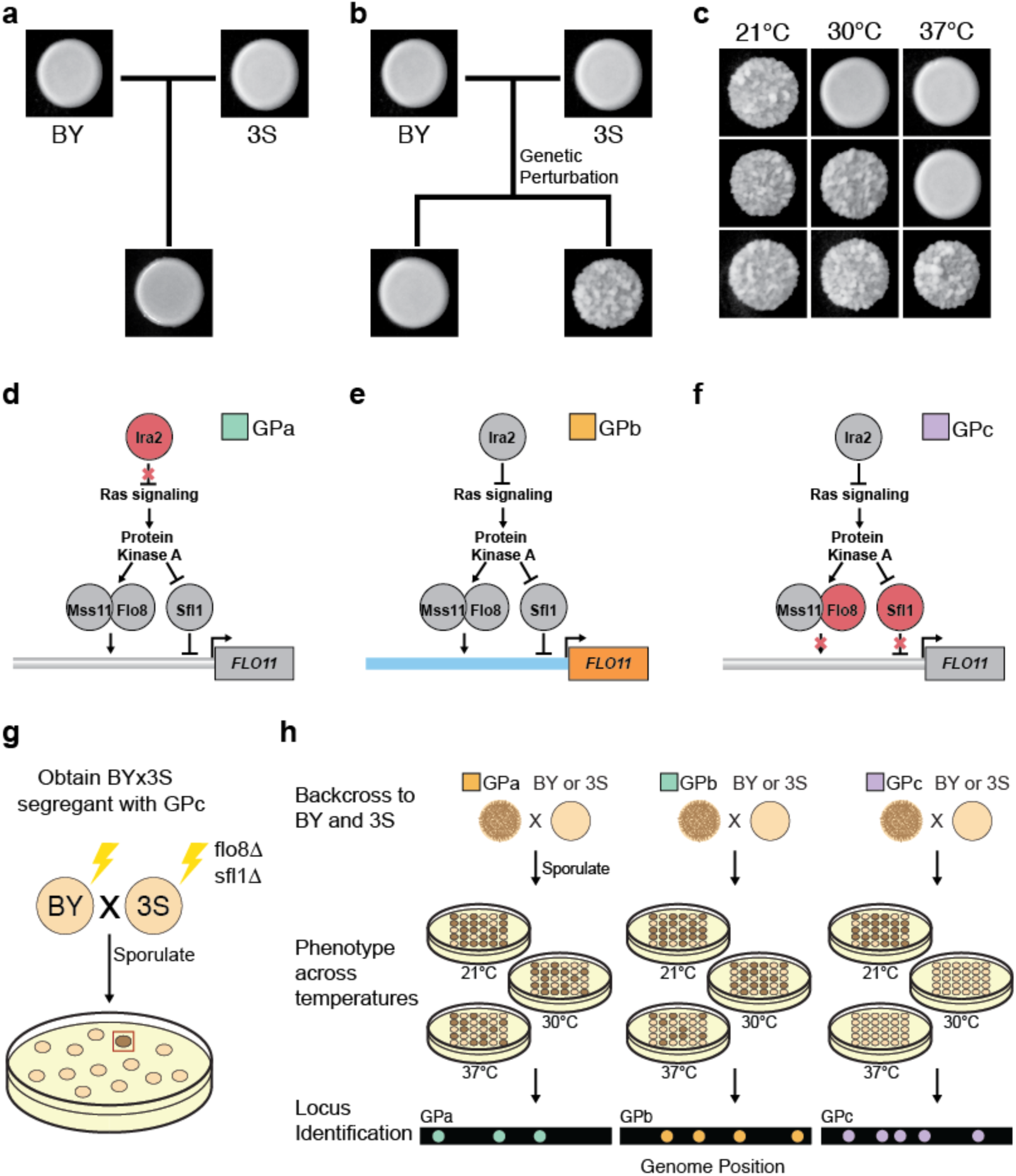
Different genetic perturbations (‘GPs’) interact with cryptic variation to cause rough morphology. **(a)** The BY and 3S strains, as well as their cross progeny, form ‘smooth’ colonies. **(b)** Certain GPs can cause BYx3S segregants to express an alternative ‘rough’ colony phenotype. **(c)** Expression of the colony trait can be temperature-sensitive, with segregating loci causing individuals form rough colonies at 21°C only, 21 and 30°C, or 21, 30, and 37°C. **(d)** GPa is a loss-of-function mutation in *IRA2*, which encodes a Ras negative regulator. **(e)** GPb results from recombination between the BY (blue) *FLO11* promoter and 3S (orange) *FLO11* coding region. **(f)** GPc is a double knockout of *FLO8* and *SFL1*, which encode the main Ras-dependent transcriptional regulators of *FLO11*. **(g)** GPc was genetically engineered in both BY and 3S, and a rough BYx3S segregant was obtained, as indicated by a brown colony in the illustration. (**h**) Genetic mapping populations were generated by backcrossing a rough segregant with each genetic perturbation to BY and 3S, and screening for the trait at 21, 30, and 37°C. Loci associated with the trait were identified in each backcross for each GP-temperature combination.

Crucially, genetic mapping and genetic engineering techniques can be used to comprehensively identify the specific GPs and cryptic variants that enable a given rough segregant to express the trait. In our initial work on colony morphology in the BYx3S cross, we found that a *de novo* loss-of-function mutation in *IRA2* (‘GPa’), a negative regulator of Ras signaling, had occurred during the generation of BYx3S segregants and enabled ~2% of cross progeny to express the phenotype (Taylor AND Ehrenreich 2014) (**Figure 1D**). Across several papers, we demonstrated that GPa causes trait expression through temperature-dependent, higher-order genetic interactions involving cryptic variants inherited from both BY and 3S (Taylor AND Ehrenreich 2014; Taylor AND Ehrenreich 2015b; Lee *et al.* 2016). Subsequently, we identified other *de novo* and induced mutations that also facilitate expression of the rough phenotype in the BYx3S cross (Taylor AND Ehrenreich 2015b; Taylor *et al.* 2016). These other mutations tended to also disrupt negative regulation of signaling and transcription within the Ras pathway, and mostly interacted with the same alleles found in the studies focused on GPa.

In a screen of 106 independently generated BYx3S crosses in which 17% of the crosses produced at least one rough segregant, we also found individuals that expressed the trait despite lacking any detectable *de novo* mutations (Taylor *et al.* 2016). Instead, two-thirds of these segregants inherited recombination events in the promoter of *FLO11* (‘GPb’, **Figure 1E**). In total, we recovered six different recombination breakpoints, which all occurred within 1.3 kb of each other. *FLO11* encodes a flocculin whose display on the cell surface facilitates cell-cell adhesion; this protein is required for expression of the rough phenotype (Lo AND Dranginis 1996; Taylor AND Ehrenreich 2015b). Genetic engineering experiments showed that the recombination events brought at least two alleles at closely linked loci in or near *FLO11* onto the same chromosome, resulting in a new *FLO11* haplotype that behaves like the mutations described in our earlier studies (Taylor *et al.* 2016). Specifically, segregants with a BY promoter and a 3S coding region had the potential to express the trait. Despite discovering GPb in this past study, we neither resolved the cryptic variant(s) in the *FLO11* promoter nor comprehensively mapped the loci enabling this GP to exert a phenotypic effect. Thus, it was not possible to compare the genetic basis of the rough phenotype in the presence of GPb to our initial work on GPa.

Here, we use the rough colony system to determine how different types of GPs interact with distinct cryptic variants in the BYx3S cross to produce the same trait. In addition to GPa and GPb, we also examine a third GP that facilitates expression of the rough phenotype. To generate ‘GPc,’ we knocked out the activator Flo8 and the repressor Sfl1 (**Figure 1G**), which are the main transcription factors that act downstream of the Ras pathway to regulate colony morphology in the cross. Previously, we showed that deletion of *SFL1* is sufficient to enable Ras-dependent cryptic variants to express *FLO11* and that transcription of *FLO11* in these *sfl1∆* segregants is Flo8-dependent (Taylor AND Ehrenreich 2015b). By eliminating these key regulators and screening for segregants expressing the trait, we sought to uncover previously unidentified cryptic variants that can also give rise to the rough phenotype. We successfully recovered one rough BYx3S *flo8∆ sfl1∆* segregant, making it possible to examine the genetic basis of the phenotype in the absence of its primary transcriptional regulators.

In this paper, we comprehensively determine the genetic basis of the rough phenotype across temperatures for GPb and GPc, and compare these results to our past work on GPa. Across the three GPs, we identify 21 loci that contribute to the rough phenotype. Of these loci, 20 show conditional phenotypic effects that are influenced by particular combinations of GP, other loci, and temperature. Although all three factors prove important, GP is by far the strongest determinant of which loci show phenotypic effects, impacting nearly all identified loci. In fact, 12 of the detected loci only exert phenotypic effects in the presence of a single GP. Additionally, we find that the identified loci exhibit varying degrees of additivity, epistasis, and genotype-environment interaction depending on the GP that uncovers them. At the molecular level, most, if not all, of the identified loci influence *FLO11* regulation in *cis* or *trans*, suggesting that our findings result from complex genetic and environmental effects on the regulation of a single key gene, *FLO11*. These findings enhance our understanding of both the extent of cryptic variation within populations and the mechanisms by which GPs reveal cryptic variants. Further, they show how conditional uncovering of cryptic genetic variation can result in highly divergent genetic architectures that produce the same trait, even within a single population examined in a common environment.

## Materials and Methods

### *Knockout of FLO8* and *SFL1 in BY and 3S*

*FLO8* and *SFL1* targeting guide RNA (gRNA) sequences were cloned into the pML104 plasmid vector, which carries Cas9 and *URA3* (Laughery *et al.* 2015). Each resulting plasmid was transformed into BY or 3S alongside a double-stranded oligo repair template for homologous recombination. The *flo8 sfl1∆* BY and 3S strains were both generated through a series of two sequential knockouts. The lithium acetate method (Gietz AND Woods 2002) was used for transformation. Oligos were 90 bases long with homology to either *FLO8* or *SFL1*, with inclusion of a stop codon followed by single base frameshift deletion in the middle. Stop codons were introduced at amino acids 155 and 39 in *FLO8* and *SFL1*, respectively. Transformed cells were plated on solid yeast nitrogen base (YNB) media lacking uracil to select for retention of the pML104 plasmid. Plasmids were then eliminated from the transformants by plating on 5-fluoroorotic acid (5-FOA). Presence of the intended genetic modifications was checked in the transformants using Sanger sequencing.

### Isolation of rough BYx3S F_2_ segregants

GPa and GPb were identified in BYx3S crosses produced by sporulating independently generated BY/3S diploids. The GPa rough F_2_ segregant included in the current study is discussed in (Taylor AND Ehrenreich 2014; Taylor AND Ehrenreich 2015b; Lee *et al.* 2016), while the GPb rough F_2_ segregant used in this study was described in (Taylor *et al.* 2016), in which it was referred to as ‘Rough segregant 13’. GPa and GPb were identified in these past studies by taking individual rough F_2_ segregants recovered from screens of BYx3S crosses, performing genetic mapping in backcross populations obtained by mating the respective rough F_2_ segregants to both BY and 3S, and analyzing whole genome sequencing data for backcross segregants. To obtain a rough F_2_ segregant with GPc, *flo8∆ sfl1∆* BY and 3S strains were mated to one another. The resulting diploid was sporulated and plated at low density (~300 colonies per plate) onto YNB agar containing canavanine to select for MAT**a** haploids using the synthetic genetic array system (Tong *et al.* 2001). After five days at 30°C, colonies were replicated onto yeast extract-peptone-ethanol (YPE) plates. A single rough colony was identified after four days of growth at 21°C.

### Generation of backcross segregants

All data for GPa was described in previous papers (Taylor AND Ehrenreich 2014; Taylor and Ehrenreich 2015b; Lee *et al.* 2016), while data for GPb and GPc were generated in this current paper. To obtain backcross progeny for genetic mapping and segregation analysis, the rough F_2_ segregant with GPb was backcrossed to wild type BY and 3S strains, while the rough F_2_ segregant with GPc was backcrossed to *flo8*∆ *sfl1*∆ BY and 3S strains. A second-generation backcross was also performed for GPb; specifically, a rough segregant from the GPbx3S cross was mated to BY. For all backcrosses, diploids were sporulated and plated at low density onto YNB plates containing canavanine to select for random MAT**a** spores using the Synthetic Genetic Array marker system (Tong *et al.*). After five days of growth at 30°C, haploid colonies were replicated onto YPE plates and incubated at 21°C. Backcross segregants expressing rough morphology after four days of growth on YPE were inoculated into liquid yeast extract-peptone-dextrose (YPD) media and grown overnight at 30°C. Freezer stocks of rough segregants were made by mixing aliquots of these cultures with 20% glycerol solution and storing these glycerol stocks at −80°C.

### Phenotyping at multiple temperatures

Cells from freezer stocks were inoculated into liquid YPD media, grown for two days at 30°C, and then pinned onto three YPE plates. Each plate was then incubated at a single, constant temperature: 21, 30, or 37°C. The colonies were then phenotyped for the colony trait after four days of growth, and designated as belonging to one of three temperature sensitivity classes: expression of the trait at 21°C only, expression of the trait at 21 and 30°C only, or expression of the trait at all examined temperatures. Genetic mapping was then performed separately on each of these temperature sensitivity classes.

### Genotyping of GPb and GPc rough segregants

Segregants from each temperature sensitivity class were inoculated into liquid YPD. DNA was extracted from overnight cultures using the Qiagen DNeasy 96 Blood and Tissue kit. Illumina sequencing libraries were then prepared using the Illumina Nextera kit, with a unique pair of dual-indexed barcodes for each individual. Between 48 and 156 segregants from each combination of GP, backcross, and temperatures sensitivity were sequenced. Sequencing was performed at the Beijing Genomics Institute on an Illumina HiSeq 4000 using paired-end 100 bp x100 bp reads. Each segregant was sequenced to an average per site coverage of at least 2.5x. Reads were aligned to either a BY or 3S reference genome using the Burrows-Wheeler Aligner (BWA) version 7 with options mem −t 20 (Li AND Durbin 2009). SAMtools was used to generate mpileup files (Li *et al.* 2009), and genome-wide allele frequencies were calculated at 36,756 SNPs that were previously identified between BY and 3S (Taylor *et al.* 2016).

### Genetic mapping of loci underlying the rough phenotype

Segregants’ genotypes at each SNP were determined using a Hidden Markov model, implemented in the HMM package in R. Loci associated with the trait for each combination of GP, backcross, and temperature sensitivity were identified using binomial tests. Sites were considered statistically significant at a Bonferroni-corrected threshold of p < .01. Multiple-testing correction was performed on each backcross population on its own, as the number of unique tests varied 837 to 1,526 among these populations. We defined the interval surrounding identified loci by computing the −log10(*p*-value) at each linked SNP and determining the SNPs at which this statistic was 2 lower than the peak marker at a given locus. These bounds were used in fine-mapping and comparison of loci detected in different combinations of GP, backcross, and temperature sensitivity.

### Testing for genotypic heterogeneity

Observed and expected haplotype frequencies for each pair of SNPs were compared using χ^2^ tests in a custom Python script. Expected haplotype frequencies were calculated as the product of the individual allele frequencies at each of the two sites. To reduce the number of statistical tests, SNPs containing the same information across all segregants in a given backcross population were collapsed into a single marker. The Benjamini-Hochberg method for False Discovery Rate (FDR) (Benjamini AND Hochberg 1995) was then implemented using the statsmodels Python module (Seabold AND Perktold 2010). A stringent FDR of 0.0001 was employed and regions of the genome within 30,000 bases of the ends of chromosomes were excluded to avoid genotyping errors.

### Exploration of additivity and epistasis

Expected multi-locus genotype frequencies were calculated as the product of the frequencies of the individual alleles segregating in a given backcross. For each population, observed and expected genotype frequencies were compared using a χ^2^ test with degrees of freedom equal to one less than the number of possible genotypes. These analyses only included loci where both the BY and 3S allele were present among the rough segregants obtained from a population, and were only performed on backcross populations in which more than one multi-locus genotype was present.

### Genetic engineering at the FLO11 promoter and other loci

Gene deletions were generated using the *kanMX* cassette (Wach *et al.* 1994) and lithium acetate transformation (Gietz AND Woods 2002). A genomic region of interest was replaced with a PCR amplicon of the *kanMX* cassette that, on each end, had 60 bases of homology to the targeted gene. Transformed cells were plated on YPD + G418 to select for integration of *kanMX* and insertion was verified using PCR.

Allele replacements were performed using a two-step, CRISPR/Cas9-mediated approach. A *kanMX* deletion strain, generated as described above, was transformed with the pML104 plasmid (Laughery *et al.* 2015) carrying the Cas9 gene and a gRNA sequence targeting *kanMX*, along with a PCR product repair template that should result in replacement of *kanMX* with the desired allele. Cells were plated on YNB plates lacking uracil to select for retention of the plasmid and then plated again onto 5-FOA to eliminate the plasmid. Replacement of the *kanMX* cassette was verified by PCR and Sanger sequencing, and transformants were plated and screened on YPE to determine the phenotypic effects of allele replacement. Also, in parallel with each allele replacement, we generated control strains where *kanMX* was replaced with the original sequence at a given site. The phenotypes of allele replacement strains were then compared to the phenotypes of these control strains that were generated in parallel. Engineering of the *FLO11* promoter was performed in the BYx3S F_2_ segregant with GPb. Gene deletions and allele replacements for other genes were performed in a representative BY or 3S backcross segregant harboring either GPb or GPc.

## Results

### Isolation of a rough flo8∆ sfl1∆ segregant

Our past work showed that expression of the rough phenotype in the BYx3S cross occurs due to genetic interactions between GPs that were not present in the cross parents and cryptic variants that segregate in the cross and mainly reside in the Ras pathway (Taylor AND Ehrenreich 2014; Taylor AND Ehrenreich 2015b; Lee *et al.* 2016; Taylor *et al.* 2016). To examine whether genetic variation beyond the Ras pathway might also be able to contribute to the trait, we knocked out *FLO8* and *SFL1* in both BY and 3S using CRISPR/Cas9 (**Methods**). We then employed random spore techniques to generate and screen > 100,000 *flo8*∆ *sfl1*∆ (GPc) cross progeny for the trait at 21°C (**Methods**). We used this condition because the trait is less genetically complex at 21°C than at higher temperatures, making it easier to find rough segregants in a screen (Lee *et al.* 2016). The GPc screen produced a single rough segregant, implying that expression of the rough phenotype in the absence of Flo8 and Sfl1 requires multiple alleles that segregate in the cross, some of which must be inherited from BY and others of which must be inherited from 3S.

### The three GPs vary in their potential to express the phenotype across temperatures

In our previous work, we showed that GPa segregants typically express the trait in a temperature sensitive manner, but that cryptic variation in the cross can eliminate this temperature sensitivity for some individuals (Lee *et al.* 2016). Because our GPb and GPc rough segregants were both obtained at 21°C, we assessed whether GPb and GPc strains also have the potential to express the trait at higher temperatures. To check this, we backcrossed GPb and GPc F_2_ segregants to both BY and 3S, and then phenotyped the resulting progeny at 21, 30, and 37°C (**Figure 1H**). Note, such backcrossing generates new multi-locus genotypes that may have the potential to express the rough phenotype at higher temperatures than the F_2_ segregants recovered from initial screens.

Upon examining 768 GPb and GPc backcross progeny at higher temperatures, we found that some GPb backcross segregants expressed the trait at higher temperatures whereas GPc backcross segregants did not (**Figure S1**). We observed varying degrees of temperature sensitivity among the rough GPb segregants, with only a minority of rough individuals capable of expressing the trait at all examined temperatures (**Figure S1**). In our past work on GPa (Lee *et al.* 2016), we also found that robustness to temperature was less frequent than temperature sensitive trait expression. This was because ability to robustly express the rough phenotype across temperatures involved more loci than temperature sensitive trait expression. Thus, when considered in light of our past findings, our current results suggest that, in the presence of GPb, ability to express the rough phenotype across temperatures is also more genetically complex than ability to express the trait only at lower temperatures. These findings also show that despite making it possible for the rough phenotype to be expressed at 21°C, GPc provides an inherently limited potential for trait expression at higher temperatures. This may be because Flo8 and Sfl1 are necessary for expression of the rough phenotype at 30 and 37°C.

### The GPs uncover distinct loci

To map loci involved in expression of the rough phenotype across temperatures, we generated > 60,000 and > 12,000 GPb and GPc backcross segregants, respectively. First-generation backcross segregants were used exclusively in mapping, except in the case of the 3S backcross of GPb at high temperature. Because expression of the rough phenotype at 30 and 37°C is rare among GPbx3S backcross segregants (**Figure S1**), determining the genetic basis of the trait among GPbx3S backcross segregants required combining information from the first-generation backcross and a second-generation backcross. Specifically, a rough first-generation GPbx3S backcross segregant was mated to 3S (**Figure S2**), which increased the frequency of the trait among backcross progeny by halving the proportion of the genome that was segregating. GPb segregants were phenotyped at 21, 30 and 37°C, while GPc segregants were only phenotyped at 21°C. Low-coverage whole genome sequencing was performed on between 48 and 151 individuals for each GP–backcross–temperature combination, and loci associated with the trait were identified based on their enrichment among genotyped segregants (**Methods**).

In past work (Figure 5 in (Lee *et al.*)), we identified eight loci that act in different multi-locus genotypes to enable GPa segregants to express the trait at particular temperatures (**Figure 2**). BY and 3S contributed the causal alleles at two and four of these loci, respectively, while the remaining two loci were detected in both the BY and 3S allele states in different multi-locus genotype and temperature contexts (**Figure 3**). Here, genetic mapping focused on GPb segregants detected a total of 11 loci across the three environments (**Figure 2**, **Figure S3**, **Table S1**). Among these loci, six and three were detected in the BY and 3S allele states, respectively (**Table 1**). The other two loci were identified in both the BY and 3S allele states, depending on the multi-locus genotype context in which they occurred. Eight of these loci influenced the trait’s expression independent of temperature. Of the remaining loci, two were detected among individuals that could express the trait at up to 30°C, while the third was identified among individuals that could express the trait at up to 37°C. In addition, we identified 12 loci in the presence of GPc (**Figure 2**, **Figure S3**, **Table S1**). Among the loci found in the presence of GPc, seven were contributed by BY and five were contributed by 3S (**Table 1**).

**Figure 2:**
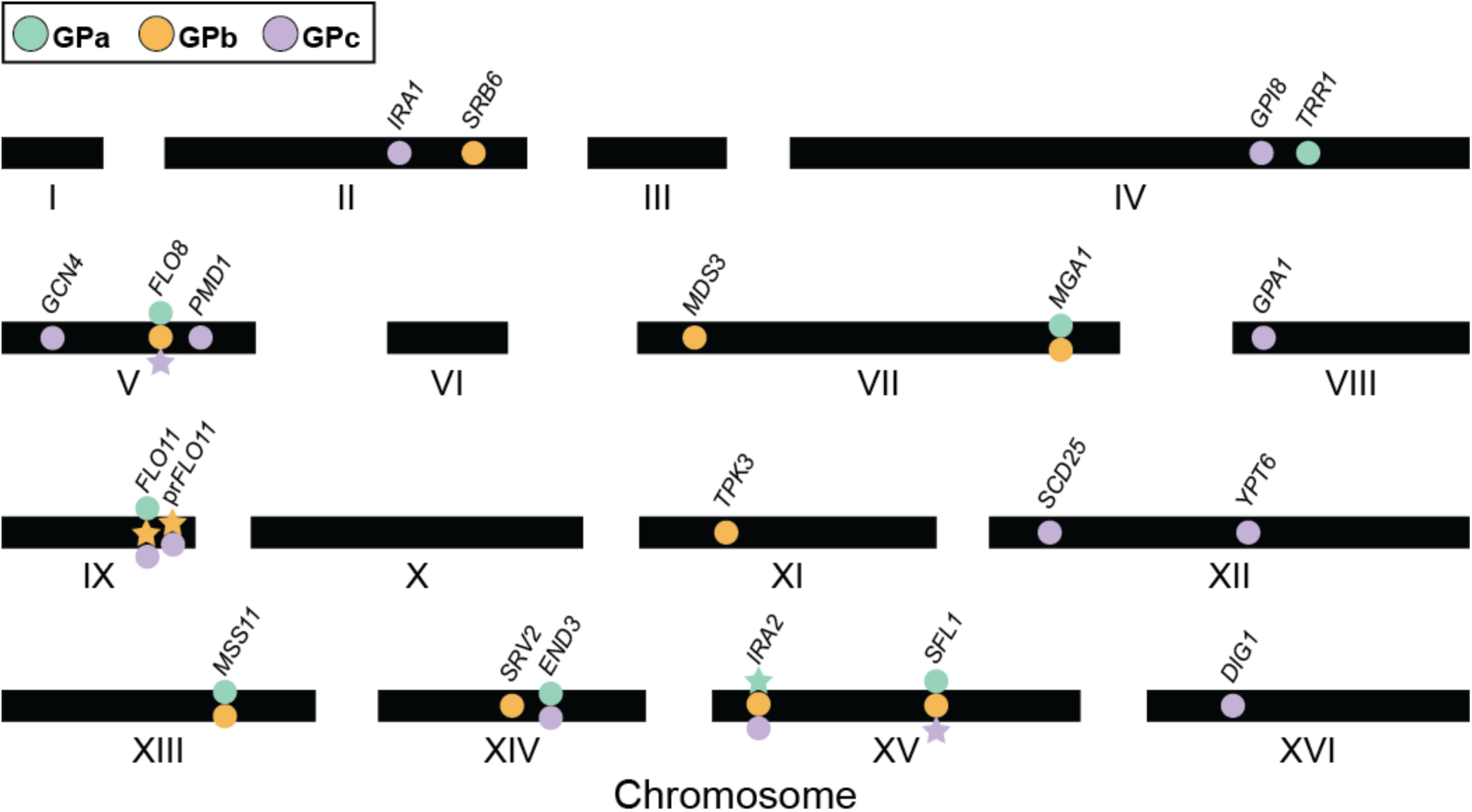
Loci identified in the presence of the GPs. Twenty-one loci were identified in total. Circles indicate cryptic variants, whereas stars indicate GPs. In some instances, the same gene that possesses a GP in one population harbors cryptic variation in another population.

**Figure 3:**
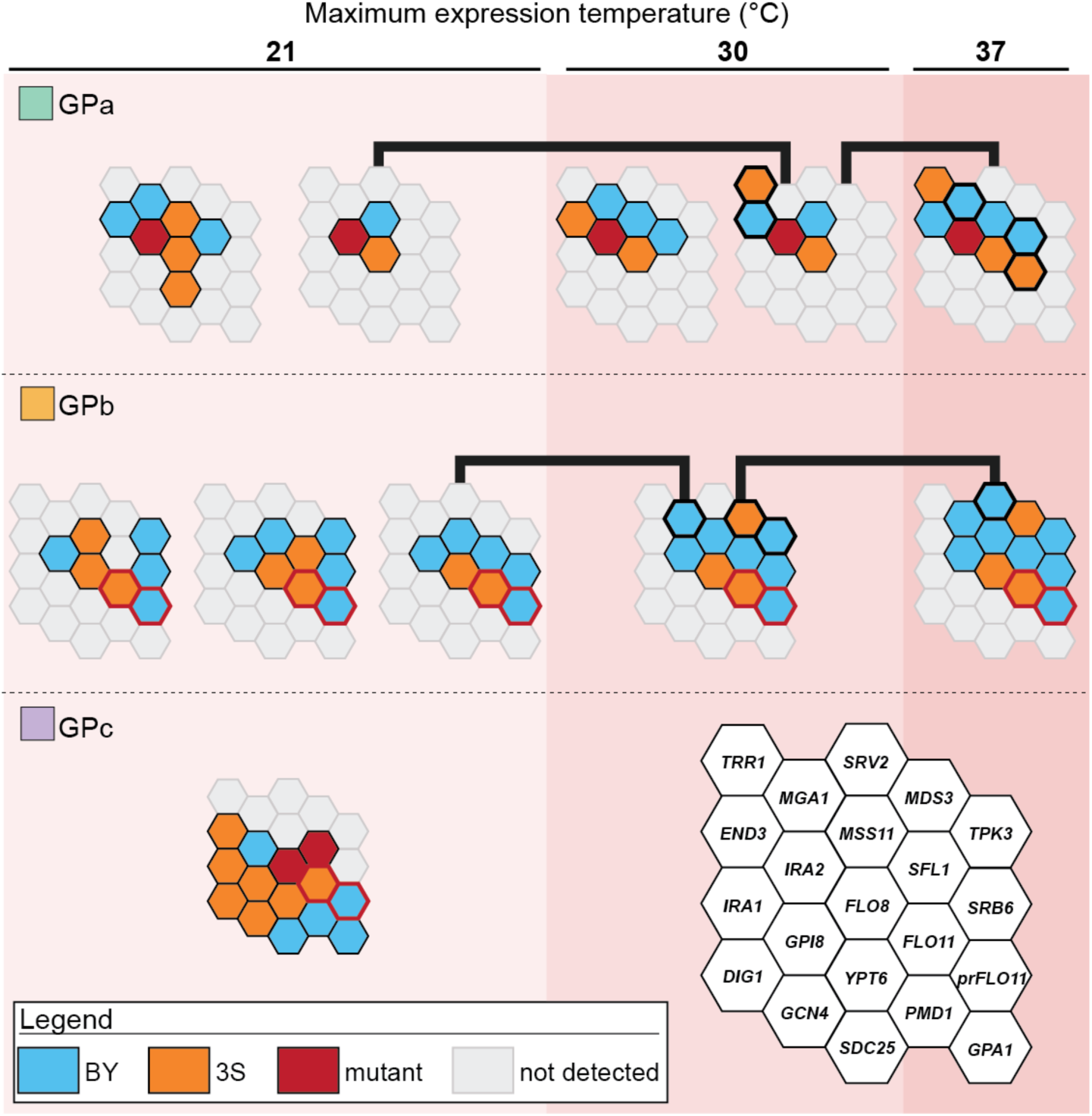
Loci detected across GPs and temperature sensitivity classes. (**a**) Loci interacting with GPa and GPb at 21 and 30°C act in different multi-locus genotypes. As shown with black lines, certain temperature-sensitive genotypes provide the potential for additional alleles to enable trait expression at higher temperatures. These additional alleles are designated with bold outlines. In the legend, ‘mutant’ refers to the <u>*de novo*</u> or induced mutations present involved in each GP. Additionally, the *prFLO11* recombination (GPb) is indicated with a red outline.

**Table 1:** Candidate genes and cloned causal genes underlying identified loci.

| Genetic Perturbation | Chr. | Gene | Function | Allele | Initial detection |
| --- | --- | --- | --- | --- | --- |
| GPc | 2 | *IRA1* | Ras negative regulator, *IRA2* paralog | 3S | Taylor et al., 2016 |
| GPb | 2 | *SRB6\** | RNA Pol II subunit | BY | this paper |
| GPc | 4 | *GPI8* | GPI transamidase complex subunit | 3S | this paper |
| GPa | 4 | *TRR1* | redox state regulator | 3S | Taylor and Ehrenreich, 2014 |
| GPc | 5 | *GCN4* | stress response transcription factor | 3S | this paper |
| GPa,b | 5 | *FLO8* | Ras activated transcription factor | 3S | Taylor and Ehrenreich, 2014 |
| GPc | 5 | *PMD1* | growth regulator; *MDS3* paralog | BY | this paper |
| GPb | 7 | *MDS3* | TOR pathway growth regulator | 3S | Taylor et al., 2016 |
| GPa,b | 7 | *MGA1* | transcriptional activator | BY | Taylor and Ehrenreich, 2015 |
| GPc | 8 | *GPA1* | G-protein | BY | this paper |
| GPa,b,c | 9 | *FLO11* | cell surface glycoprotein | 3S | Lee et al., 2016 |
| GPb,c | 9 | *prFLO11* | *FLO11* promoter | BY | Taylor et al., 2016 |
| GPb | 11 | *TPK3* | Protein Kinase A subunit | BY | Taylor et al., 2016 |
| GPc | 12 | *SDC25* | Ras GEF | BY | this paper |
| GPa,c | 12 | *YPT6* | Rab GTPase | 3S | Lee et al., 2016 |
| GPa,b | 13 | *MSS11* | Ras activated transcription factor | BY,3S | Taylor and Ehrenreich, 2014 |
| GPb | 14 | *SRV2* | activator of adenylate cyclase | BY | this paper |
| GPa,c | 14 | *END3* | cell wall morphogenesis | BY, 3S | Taylor and Ehrenreich, 2014 |
| GPb,c | 15 | *IRA2* | Ras negative regulator | BY | Taylor et al., 2016 |
| GPa,b | 15 | *SFL1* | Ras inactivated transcription factor | BY | Taylor and Ehrenreich, 2015 |
| GPc | 16 | *DIG1* | MAPK-regulated transcription factor | 3S | this paper |

\* candidate gene

Some of the loci detected across GPs and temperatures both overlapped and showed involvement of the same parental allele, indicating potential involvement of the same underlying cryptic variants. Supporting this possibility, each locus also corresponded to a causal gene that we had cloned during our work on GPa (Taylor and Ehrenreich; Taylor and Ehrenreich; Lee *et al.*; Taylor *et al.*). Assuming that these overlapping regions represent the same cryptic variant, we consolidated all detected genomic intervals into 21 distinct loci (**Figure 2**, **Table 1**). Among these loci, one was found in the presence of all three GPs, 8 were found in the presence of two GPs, and 12 were found in the presence of a single GP. This indicates that nearly all of the detected loci show differential responsiveness to the GPs and that the majority of the loci we have identified are cryptic variants that only act in the presence of specific GPs. However, if our assumption of overlapping loci representing the same cryptic variants is incorrect, then the responsiveness of the identified cryptic variants to GP and temperature would be even higher than discussed in this paper.

### The GPs alter genotype-environment interaction

We assessed how identified loci interact with particular GPs, each other, and temperature to produce the rough phenotype. Given that we performed such an analysis on GPa in the past (Lee *et al.* 2016) and GPc segregants can only express the trait at 21°C, we focused this analysis on GPb. Note, our past work on GPa found that at particular levels of temperature sensitivity, distinct sets of epistatic cryptic variants form multi-locus genotypes that produce the phenotype (Lee *et al.* 2016). For example, at 30°C, two distinct combinations of four and five cryptic variants act in conjunction with GPa to cause the trait’s expression (**Figure 3**). Examination of first-generation GPb backcross segregants produced results comparable to our findings for GPa. Among GPb segregants expressing the trait exclusively at 21°C, we detected a pair of interacting loci on chromosomes XIII and XV, which then allowed us to identify distinct multi-locus genotypes present among these individuals (**Figure 3**, **Figure S4**, **Note S1**, **Methods**). Similar to our work on GPa (Lee *et al.* 2016), only one of the GPb multi-locus genotypes found at 21°C provided a foundation upon which additional allele substitutions can facilitate trait expression at higher temperatures (**Figure 3**, black lines). Among these GPb segregants exhibiting the trait at higher temperatures, single multi-locus genotypes indicative of higher-order epistasis facilitated trait expression at temperatures up to 30 and 37°C (**Figure 3**). Comparison our results for GPb to our past work on GPa found that more than half of the loci and the exact multi-locus genotypes differed between the two GPs (**Figure 3**). These results show that the GPs significantly modify genotype-environment interactions underlying the rough phenotype.

### The GPs affect additivity and epistasis among identified loci

The only temperature at which all three GPs enable trait expression is 21°C. To compare the quantitative genetic architectures enabling the three GPs to induce the rough phenotype at this temperature, we used genotype data from backcrosses (**Figure 4A** through **C**, **Methods**). If loci act in a predominantly additive manner in the presence of a given GP, observed multi-locus genotype frequencies should match expected multi-locus genotype frequencies, which can be calculated as the product of the frequencies of every individual allele involved in a multi-locus genotype. In contrast, if epistasis meaningfully contributes to the trait, observed multi-locus genotype frequencies should depart from expected multi-locus genotype frequencies. Based on these tests, we observed significant deviation from expected frequencies for GPa (**Figure 4A**) and GPb (**Figure 4B**), confirming that epistasis plays a significant role in the trait in the presence of these GPs. However, data for GPc suggested that trait expression in the presence of this GP is entirely additive (**Figure 4C**). These findings imply that although the three GPs in this study can each enable expression of the same rough phenotype, they do so not only through largely distinct cryptic variants but also through fundamentally different quantitative genetic architectures.

**Figure 4:**
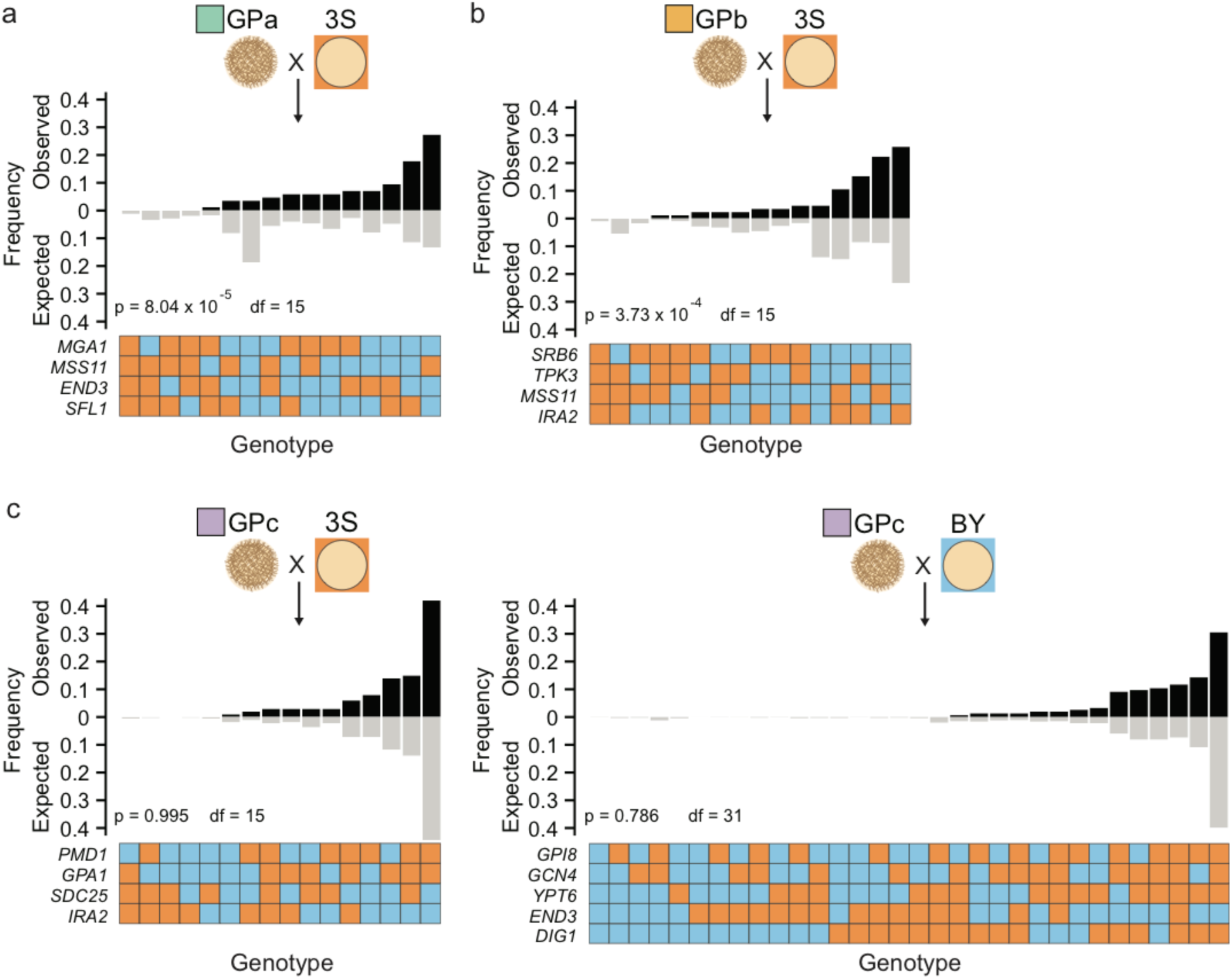
Loci exhibit epistasis or additivity depending on GP. (**a**) For GPa individuals expressing the trait at only 21°C, observed multi-locus genotype frequencies at the four loci segregating in the 3S backcross do not reflect expected frequencies. This can occur if certain alleles exhibit epistasis and co-segregate in a non-random manner. **(b)** Observed and expected genotype frequencies based on four loci in GPb individuals expressing the trait at only 21°C. **(c)** Observed genotype frequencies in GPc individuals closely match expected frequencies, suggesting involved loci act in an additive manner. In all cases, expected genotype frequencies are computed as the product of the frequency of the involved alleles. P-values correspond to results of a χ^2^ test with degrees of freedom indicated in the figure. Loci for which only a single allele is present in their respective population were excluded from analysis, and backcross populations in which only one multi-locus genotype was present are not shown. Only first-generation backcross segregants were used to generate this figure.

### Coding and regulatory variation in FLO11 plays an essential role in the trait

Only a single locus exhibited a phenotypic effect in the presence of all three GPs. This locus corresponds to the 3S allele of the *FLO11* coding region (Lee *et al.* 2016). The BY and 3S alleles of *FLO11* possess not only 57 synonymous and 29 nonsynonymous SNP differences, but also a length polymorphism of ~581 nucleotides (**Figure S5)**. *FLO11*^3S^ encodes a longer cell-surface protein than *FLO11*^BY^, an attribute that is known to favor the expression of multi-cellular traits like the rough phenotype (Verstrepen *et al.* 2005; Fidalgo *et al.* 2006; Fidalgo *et al.* 2008; Zara *et al.* 2009; Hope AND Dunham 2014; Matsui *et al.* 2015). Although this length polymorphism is likely causal for the trait, we cannot rule out the possibility that some of the SNPs also play a role.

In addition to being a required component of GPb, we found the BY allele of the *FLO11* promoter is necessary for GPc segregants to express the trait (**Figure 3**, **Table 1**). *FLO11* has one of the largest and most complex promoters in *S. cerevisiae*, with more than 17 transcription factors and 6 signaling cascades capable of influencing its regulation (Bruckner AND Mosch 2012). Through genetic engineering experiments, we localized the causal variant in the *FLO11* promoter to a Rim101 binding site that is present in 3S but not BY (**Figure 5)**. This finding is consistent with the important role that transcriptional derepression of *FLO11* plays in expression of the rough phenotype (Taylor AND Ehrenreich 2015b; Taylor *et al.* 2016). We note that although Rim101 has been described as a *FLO11* activator in other strains of *S. cerevisiae*, this role is indirect and mediated through its role in silencing *NRG1* (Kuchin *et al.* 2002; Lamb AND Mitchell 2003; Barrales *et al.* 2008), which encodes a repressor that directly binds the *FLO11* promoter. The Rim101 binding site in the 3S *FLO11* promoter most likely results in direct repression of *FLO11* by Rim101, which reinforces Sfl1-mediated repression. These findings not only speak to the critical role of Flo11 in expression of the rough phenotype in the BYx3S cross, but also illustrate how regulation of this gene by multiple pathways determines the phenotypic effects of cryptic variation.

**Figure 5:**
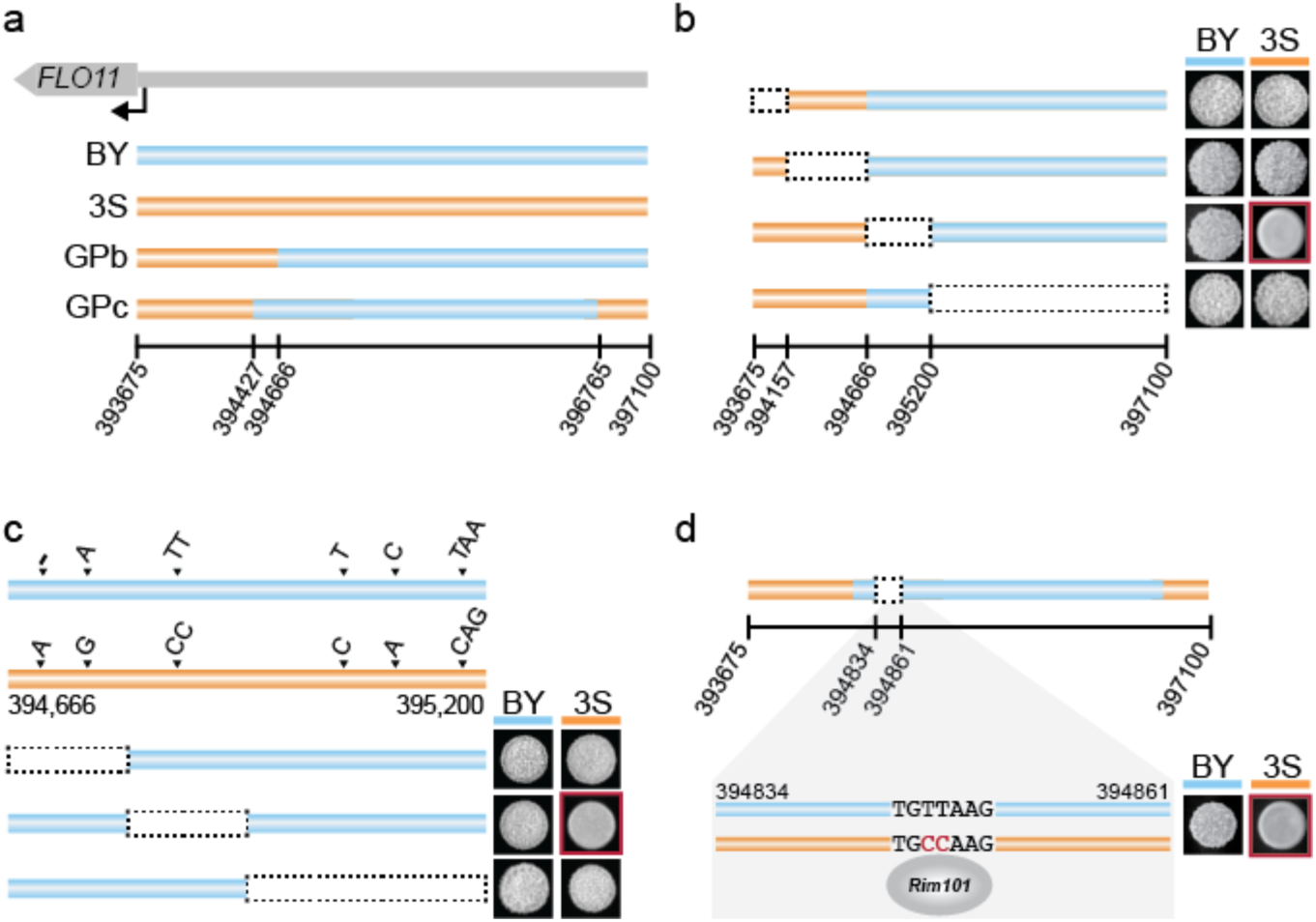
A transcription factor binding site polymorphism is required for GPb and GPc rough segregants to express the trait. (**a**) The GPb and GPc *FLO11* promoters contain portions of both the BY (blue) and 3S (orange) versions. **(b)** Genetic engineering experiments identified a 534 nucleotide segment of the BY-derived promoter as being responsible for the trait. The causal replacement with the 3S allele leading to loss of the phenotype is outlined in red. **(c)** Engineering of the six polymorphisms within the delimited promoter region revealed that replacement of the TT BY sequence with the 3S CC at position 394,846 to 394,847 allele ablates the phenotype. **(d)** The causal polymorphism results in a Rim101 binding site only present on the 3S promoter. In panels **b** through **d**, red boxes denote allele replacements with phenotypic effects.

### GPa and GPb rough segregants utilize different sub-pathways that impact Ras

Excluding the *FLO11* coding region and promoter, all remaining loci act in the presence of only one or two of the GPs. To better understand the highly contextual effects of these cryptic variants, we attempted to resolve detected loci to specific genes. Seven of the GPb-responsive loci detected in this study were previously found to interact with either GPa or other GPs in previous papers, in which they were mapped to individual genes (Lee *et al.* 2016; Taylor *et al.* 2016) (**Table 1**). These genes encode Ras-regulated transcription factors (*FLO8*, *MSS11*, *MGA1*, *SFL1*), a protein kinase A subunit (*TPK3*), a TOR pathway component (*MDS3)*, and *IRA2*. Two of the loci found in the current study among GPb segregants were located on chromosomes II and XIV, and had never been detected in our past work (**Figure 2, Table 1**). Through genetic engineering experiments, we resolved the chromosome XIV locus to *SRV2* (**Figure S6)**, which encodes a post-translational activator of adenylate cyclase, another component of the Ras pathway. At the chromosome II locus, the most likely candidate is *SRB6*, which encodes an essential subunit of the RNA polymerase II mediator complex. Previously, we showed that mutations disrupting other mediator components, Srb10 and Srb11 (also known as Ssn3 and Ssn8, respectively), can induce the rough phenotype by interacting with a subset of the alleles that have a phenotypic effect in the presence of GPa (Taylor *et al.* 2016) (**Note S2**).

Although loci interacting with GPa and GPb primarily act through the Ras pathway and many loci interact with both GPs, some are specific to one or the other. Among these are *TRR1* and *END3*, which were detected in GPa segregants but not GPb segregants, as well as *MDS3*, *SRV2*, and *TPK3*, which were identified in GPb segregants but not GPa segregants. Notably, these genes play a role in activating the Ras pathway through oxidative stress and actin organization (**Figure 6**). Actin cytoskeleton stability is required for cell polarity and yeast-adhesion traits, and is regulated in part by the effects of End3 and Srv2 on Ras signaling (Du AND Ayscough 2009). This process results in increased production of reactive oxygen species (ROS) through the activity of Tpk3 (Gourlay AND Ayscough 2006). ROS accumulation is then influenced by Trr1 (Charizanis *et al.* 1999) and the TOR pathway, of which Mds3 is a component. Together, Mds3, Srv2, and Tpk3 form a well-described sub-pathway that influences Ras activity (Gourlay AND Ayscough 2006; Du AND Ayscough 2009). Thus, although different loci cause the trait in the presence of GPa and GPb, they appear to reflect different sub-pathways affecting the same cellular processes, which ultimately impact how Ras-regulated transcription factors influence *FLO11* activity.

**Figure 6:**
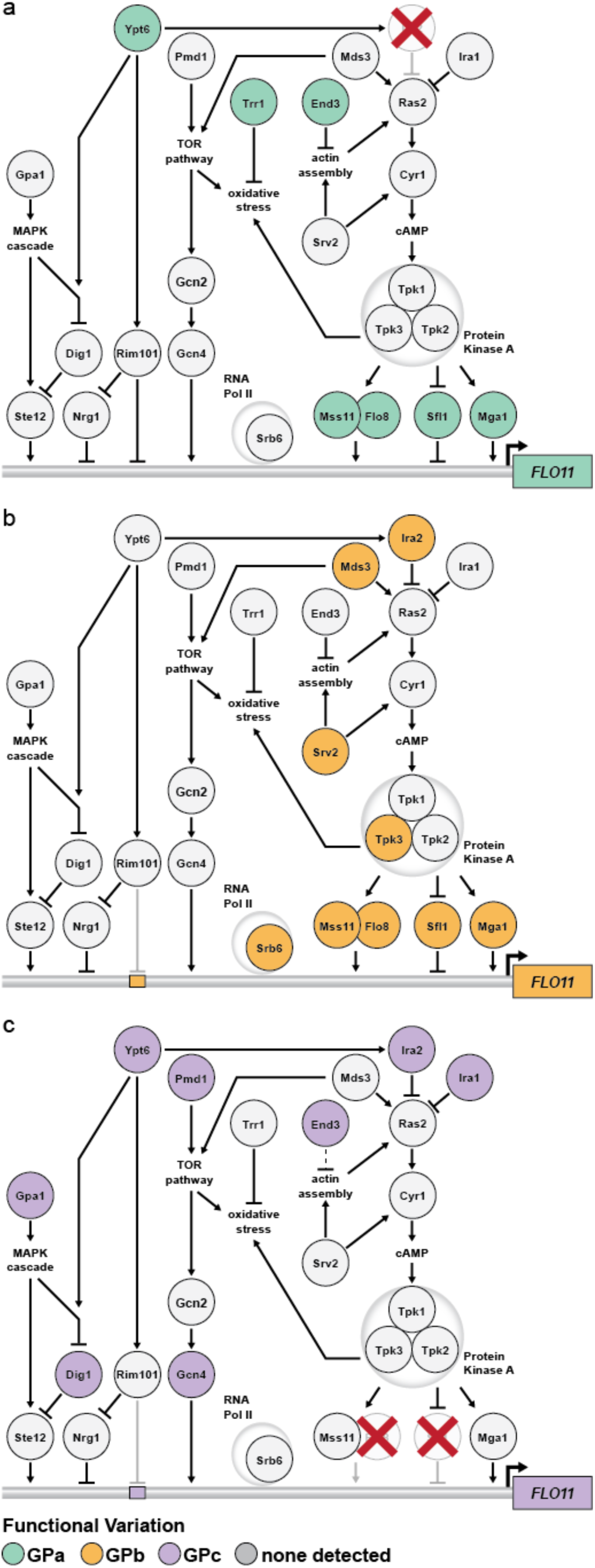
GPs unmask cryptic genetic variation in parallel signaling pathways and sub-pathways. (**a**) Loci that interact with GPa influence *FLO11* regulation by the Ras pathway. **(b)** GPb uncovers loci that primarily act through a different Ras sub-pathway involving Mds3, Srv2, and Tpk3. **(c)** Loci interacting with GPc function in a number of different pathways that are capable of regulating *FLO11* activity. The locations of transcription factor binding sites are not intended to reflect specific positioning along the *FLO11* promoter.

### Cryptic variation in several pathways underlies the phenotype in GPc segregants

Lastly, we sought to determine the genes harboring cryptic variants that interact with GPc. Regarding GPc, 3 of the 10 identified loci also interact with either GPa or b. These loci correspond to *END3*, *IRA2*, and a locus on chromosome XII. For chromosome XII and the remaining seven loci, we identified likely candidate genes based on our highly resolved genetic mapping data and publicly available research on these genes’ functions and phenotypic effects. One of these loci corresponds to *IRA1*, a Ras negative regulator and paralog of *IRA2* (Lee *et al.* 2016) (**Table 1**). To obtain additional support for the remaining loci, we performed gene deletions and allele replacements on the identified candidate genes in a GPc rough segregant, and determined the resulting effects on the colony trait (**Methods**). Knock out and replacement of *PMD1, GPA1, SDC25, GPI8, GCN4,* and *YPT6* resulted in loss of the phenotype (**Figure S7a** and **b**), implying that these six genes play positive roles in the phenotype’s regulation. In contrast, deletion of *DIG1*, a protein that directly inhibits the transcriptional activator Ste12, enhanced the trait’s expression (**Figure S7a** and **c**). These results suggest that genetic variation in these genes plays a causal role in the rough phenotype.

Detection of *END3*, *IRA1*, and *IRA2* in the absence of Flo8 and Sfl1 indicates that Ras still contributes to *FLO11* regulation in the absence of these transcription factors, possibly by impacting the activities of other transcription factors (Estruch 2000). The remaining loci implicate alternative signaling pathways as being responsible for trait expression in GPc segregants. The genes we identified in the other intervals as having a phenotypic effect when knocked out were a TOR pathway component and *MDS3* paralog (*PMD1*), members of the MAP kinase (MAPK) signaling cascade (*DIG1*, *GPA1*) (Metodiev *et al.* 2002), an environmentally-responsive transcriptional activator (*GCN4),* a Rab GTPase that influences Ras (Costanzo *et al.* 2010), MAPK (Costanzo *et al.* 2016), and Rim101 (Zheng *et al.* 2010) signaling *(YPT6),* a stress responsive guanine exchange factor (*SDC25*), and an enzyme that post-translationally modifies proteins to help them anchor into the cell wall (*GPI8*). Of particular note for *GPA1*, the BY strain is known to carry a lab-derived allele that is an expression QTL hotspot (Yvert *et al.* 2003), supporting the possibility that *GPA1* allele state might also impact expression of the rough phenotype in the presence of GPc. Additionally, *GPA1*^BY^ was previously shown to influence other *FLO11*-dependent traits through its downstream transcriptional activator Ste12 (Matsui *et al.* 2015). Nearly all of the genes implicated in allowing GPc segregants to express the rough phenotype have the potential to influence, either directly or indirectly, *FLO11* transcription. The lone exception is *GPI8*, which could still influence Flo11 at the protein level, as it is responsible for adding glycosylphosphatidylinositol (GPI) anchors to new proteins and could affect Flo11’s binding to the cell surface (Benghezal *et al.* 1996). These results show the abundant cryptic variation that impacts regulation of *FLO11* and the rough phenotype.

## Discussion

To better understand the extent of cryptic variation within a population, as well as the mechanisms regulating this cryptic variation, we comprehensively determined the genetic basis of a model phenotype that is only expressed in the presence of particular GPs. By doing this, we identified 21 loci harboring cryptic variants that can contribute to the rough phenotype. Notably, all but one of these loci show phenotypic effects that depend on the GP that is present. In addition, because the cryptic variants that are uncovered by each GP vary in their degree of epistasis with each other and interaction with the environment, we find that the trait’s genetic architecture significantly differs across the GPs. This results in a multitude of genotype, environment, and phenotype relationships that produce the same trait.

Given the detailed understanding that we have obtained for the colony morphology system, a major question is, how might our findings relate to other traits and species? One major insight from our study that may apply to other systems is that most, if not nearly all, of our findings connect to the transcriptional regulation of a single key gene, *FLO11*. Supporting this point, the single locus common to all three GPs is the coding region of *FLO11*, suggesting that regulation of Flo11 levels and stability is the central determinant of the rough phenotype’s expression across GPs, combinations of segregating loci, and temperature. Bolstering the importance of variability in *FLO11* regulation to our findings, most of the loci that exhibit phenotypic effects influence *FLO11* in *trans.* Moreover, many of these loci are only visible in the absence of a repressive Rim101 binding site in the *FLO11* promoter. These results highlight the potentially important role that *cis*-regulatory polymorphisms can play in enabling *trans*-regulatory polymorphisms to exert phenotypic effects. Indeed, *cis*-regulatory polymorphisms have been shown to modify the effects of *trans* variants in other systems (Reddy *et al.* 2012; Wong *et al.* 2017), thereby altering how genetic differences impact traits (Payne AND Wagner 2014).

In addition to showing that different GPs enable distinct cryptic variants to have phenotypic effects, we also demonstrated that the GPs modify how genotype-environment interactions impact the trait. For both GPa and GPb, we find that certain combinations of epistatic alleles both enable expression of the trait at a permissive temperature of 21°C, as well as provide the genetic potential for the phenotype at higher temperatures. However, different loci and multi-locus genotypes enable GPa and GPb segregants to express the trait at higher temperatures. In contrast, GPc individuals are unable to express the phenotype at temperatures above 21°C, implying their potential to express the trait across environments is constrained. These findings support the concept that not all genotypes specifying the same trait possess comparable phenotypic robustness (Wagner 2012; Payne *et al.* 2014; Pfennig AND Ehrenreich 2014; Siegal AND Leu 2014; Ehrenreich AND Pfennig 2016). In the case of the rough phenotype, our genetic mapping and genetic engineering results imply that differences in phenotypic robustness relate to changes in signaling and transcription factor activity across multiple pathways and sub-pathways influencing *FLO11*.

Furthermore, whereas the rough phenotype in GPa and GPb segregants involves complex epistatic effects, we find no evidence for epistasis in GPc rough segregants. This implies that by eliminating Flo8 and Sfl1, the main transcriptional regulators of *FLO11*, we not only uncovered a different set of cryptic variants but also converted the trait’s genetic architecture from mainly epistatic to additive. Perhaps this additivity at the phenotypic level reflects the cumulative effect of multiple pathways influencing *FLO11* expression at the molecular level. While the majority of the loci that interact with GPa and GPb correspond to components of the Ras pathway, many of the loci found in the presence of GPc are involved in other signaling pathways that may have compensatory functions (**Figure 6**). This is consistent with the idea that eliminating Flo8 and Sfl1 might enable other pathways to play a stronger role in expression of *FLO11* and the rough colony phenotype.

Although not an explicit focus of the current manuscript, our results also provide valuable insights into genetic background effects, the phenomenon in which GPs show different phenotypic effects in distinct individuals (Nadeau 2001; Chandler *et al.* 2013; Ehrenreich 2017). A number of recent studies have shown that these background effects often result from higher-order epistasis between a GP and multiple segregating loci (Chandler *et al.* 2014; Taylor AND Ehrenreich 2014; Miotto *et al.* 2015; Taylor AND Ehrenreich 2015b; Taylor AND Ehrenreich 2015a; Lee *et al.* 2016; Kuzmin *et al.* 2018; Mullis *et al.* 2018). Despite supporting an important role for higher-order epistasis in background effects, our current work, in particular on GPc, also shows that background effects can have much simpler underpinnings. In the context of GPc, identified loci each show pairwise epistasis with the GP, but exhibit no epistasis with each other and thus appear to act additively. These differences in quantitative genetic architecture again appear to tie back to *FLO11* regulation, consistent with theoretical work suggesting that how GPs perturb gene regulatory networks can impact whether loci show additive or epistatic effects (Gjuvsland *et al.* 2007).

In summary, our study demonstrates the large amount of cryptic variation that can underlie a single phenotype and strongly implicates complex changes in transcription as a mechanism regulating this cryptic variation. Further, these results suggest a broader insight into the genetic architecture of complex traits. Although it has long been known that traits can vary in genetic architecture depending on the different populations and environments in which they are measured, our findings illustrate that even the same phenotype examined within a single population can exhibit a spectrum of genetic architectures. As shown by the GPs in our study, which of these architectures is visible may depend on only one or two alleles that modify how the rest of the genetic variation in the population behaves. This suggests that characterizing the molecular mechanisms that shape genetic architecture will be an important step in improving our basic understanding of the relationship between genotype and phenotype.

## Acknowledgments

We thank Joseph Hale, Takeshi Matsui, Martin Mullis, Joann Phan, Rachel Schell, Fabian Seidl, Matthew Taylor, and Gleidson Teixeira for valuable input regarding this project or manuscript. We also thank the anonymous reviewers of this paper for their helpful feedback. The work reported in this paper was supported by grant R01GM110255 from the National Institutes of Health to I.M.E, a fellowship from the Alfred P. Sloan Foundation to I.M.E, and a fellowship from the São Paulo Research Foundation to A.L.V.C.

## Data Availability

All sequencing data from this project is available through the National Center for Biotechnology Information (NCBI) Short Read Archive. Data can be accessed under Bioproject ID PRJNA503265, with Biosample accessions SAMN10356503-SAMN10357319.

---

**Figure S1:**
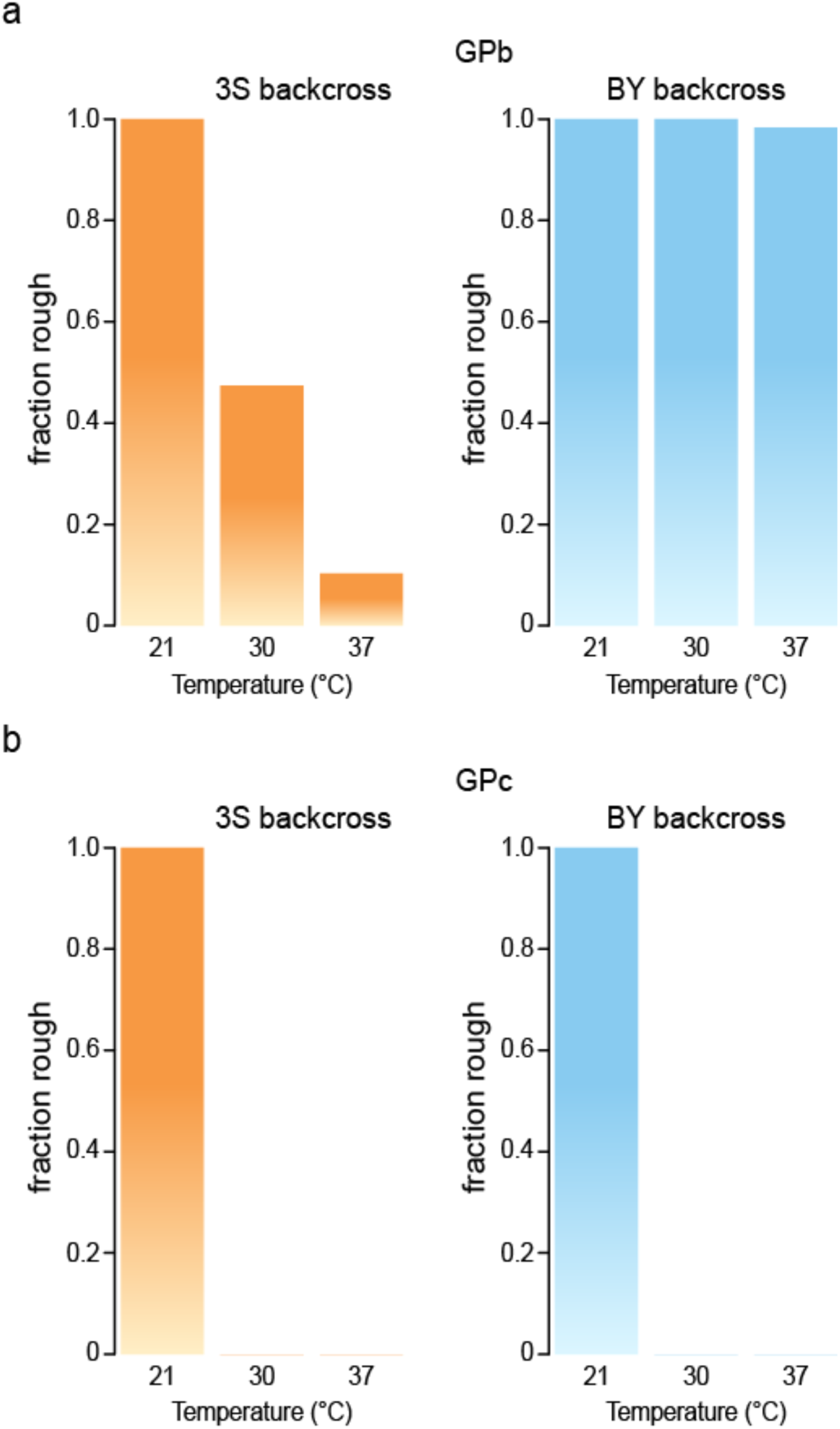
GPb and GPc individuals differ in their temperature sensitivities. **(a)** A subset of GPb backcross segregants express rough morphology at 30 and 37°C. **(b)** GPc backcross segregants are incapable of expressing the trait at 30 and 37°C. Ninety-six backcross segregants were generated and phenotyped for each GP-backcross parent combination.

**Figure S2:**
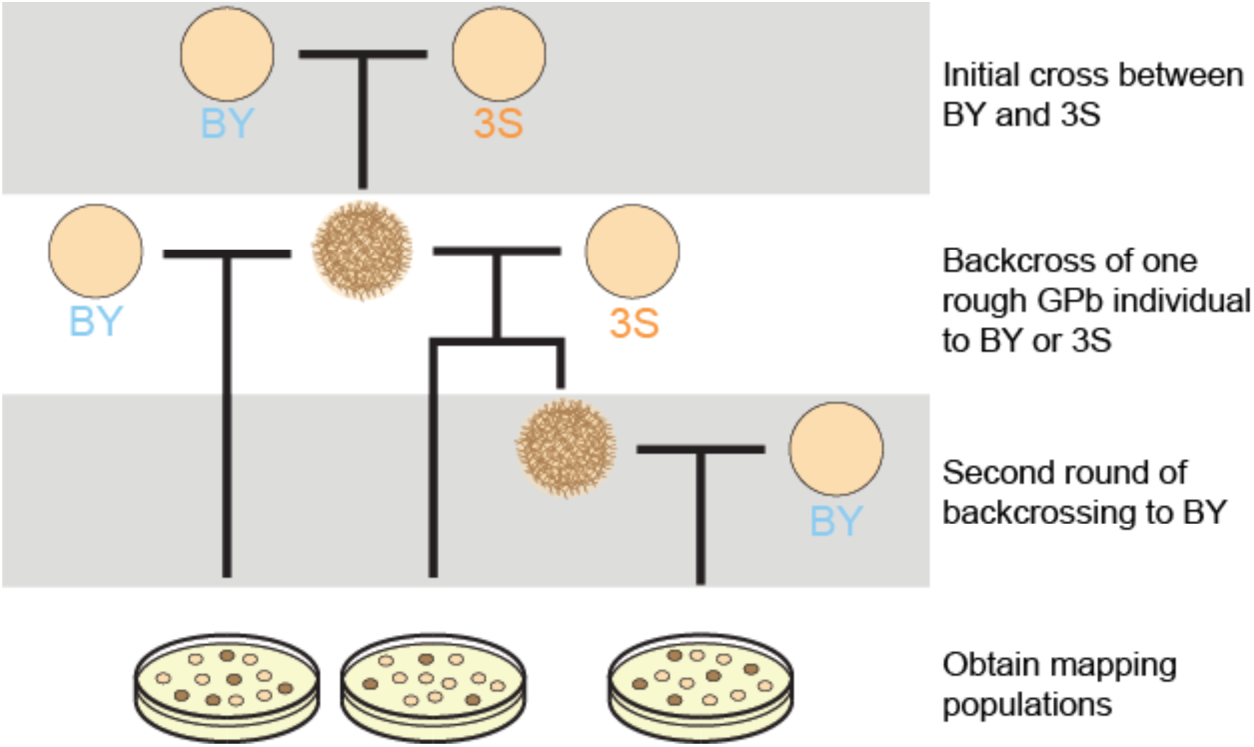
Scheme to generate GPb backcross mapping populations. A rough BYx3S F_2_ segregant carrying GPb was backcrossed to BY and 3S. In the case of the cross to 3S, a second-generation backcross to BY was then performed, which was necessary to enrich for segregants that express the rough phenotype at 37°C.

**Figure S3:**
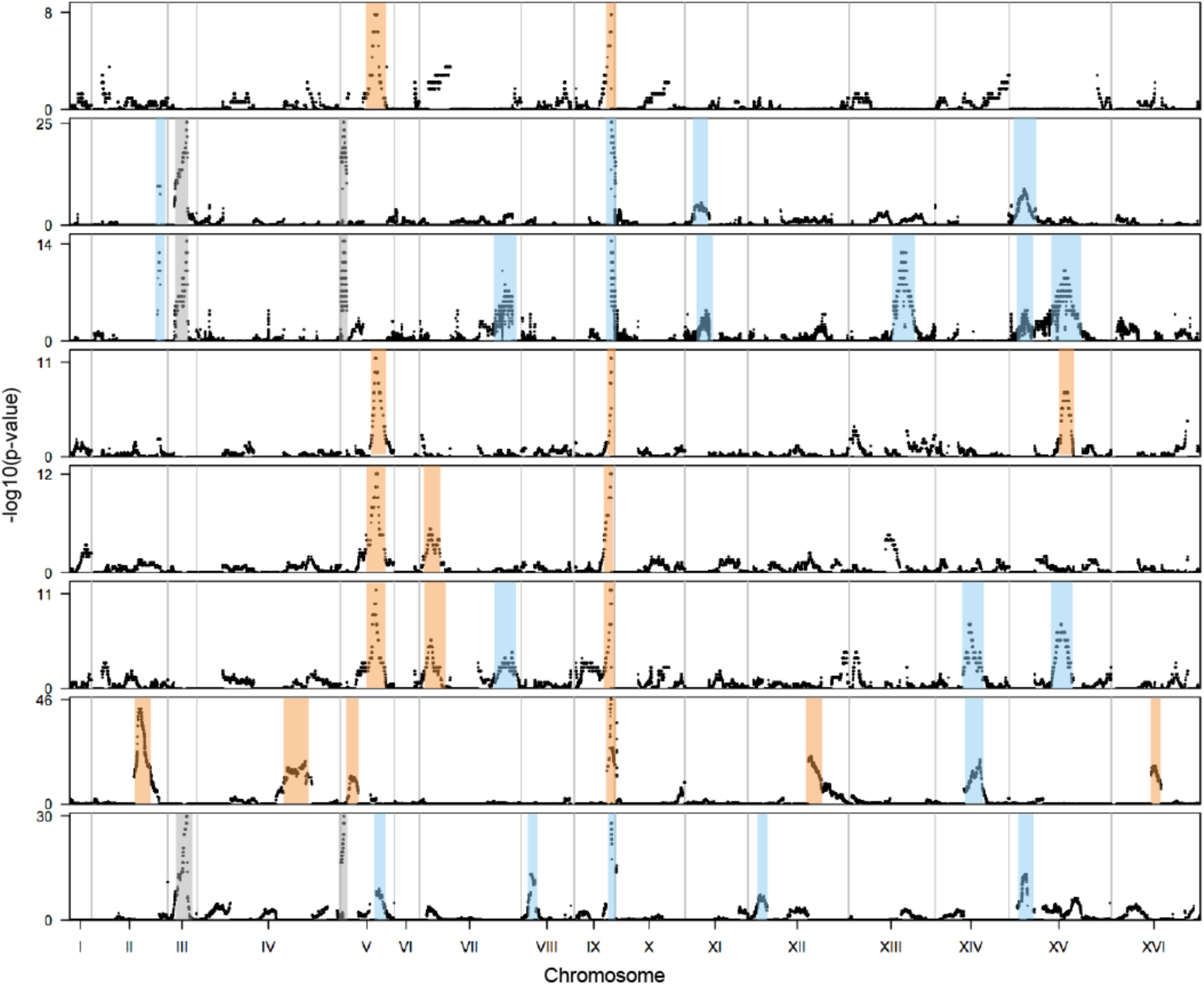
Genetic mapping for loci interacting with GPb and GPc. For each mapping population based on GP, backcross, and temperature sensitivity, a binomial test was performed for each SNP to identify regions of the genome associated with the rough phenotype. Results in are plotted in −log_10_(p)-values in the following order: GPb (BY backcross, 21, 30, 37°C), GPb (3S backcross, 21°C), GPb (3S backcross, 21, 30°C), GPb (2^nd^ generation backcross, 21°C), GPb (2^nd^ generation backcross, 21, 30°C), GPb (2^nd^ generation backcross, 21, 30, 37°C), GPc (BY backcross, 21°C), GPc (3S backcross, 21°C). Blue bars indicate loci originating from BY, and orange bars indicate loci from 3S. Gray bars denote selectable markers used in generating mapping populations. Details regarding each locus are found in Table S1.

**Figure S4:**
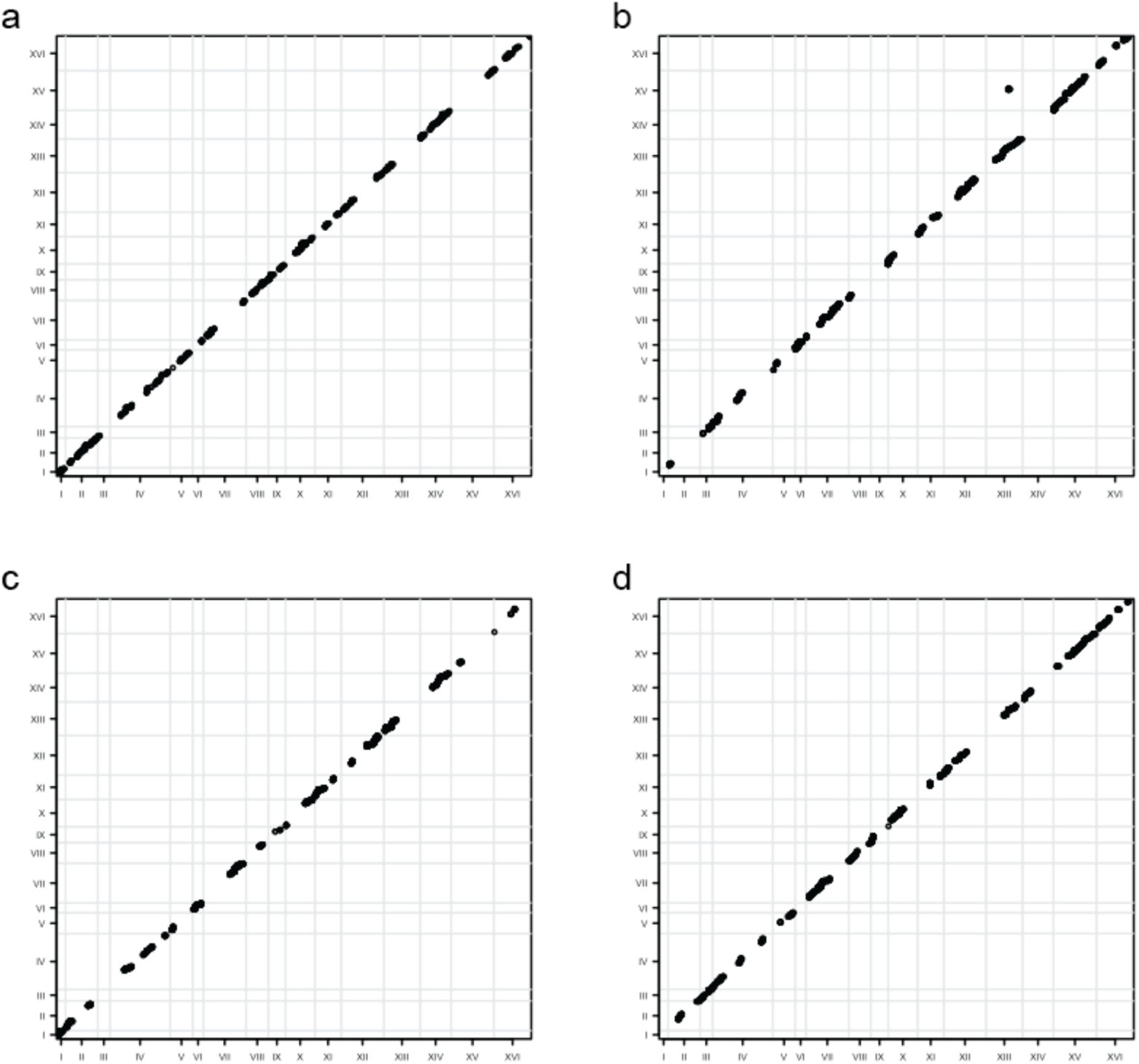
Scan for loci with correlated allele states. Significant p-values for χ^2^ tests between each pair of uniquely segregating genomic regions: GPb – **(a)** backcross to BY, **(b)** backcross to 3S, and GPc – **(c)** backcross to BY, **(d)** backcross to 3S. Results are only plotted for the upper triangle. Significant results were only found in the GPb backcross to 3S, identifying an interaction between the genes *MSS11* and *SFL1*.

**Figure S5:**
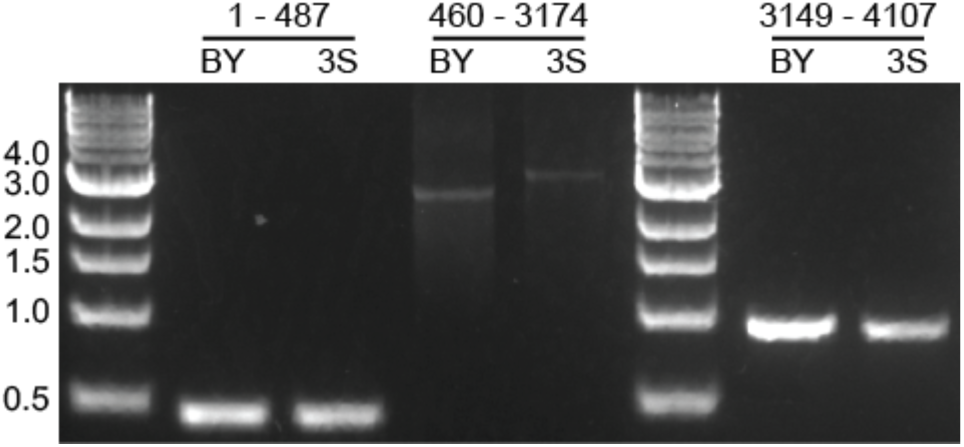
Variation in the *FLO11* coding region. The 3S strain carries a longer *FLO11* allele relative to BY. This is due to a ~500-600 nucleotide length polymorphism in the middle of the gene. Nucleotide positions are provided based on the BY gene sequence.

**Figure S6:**
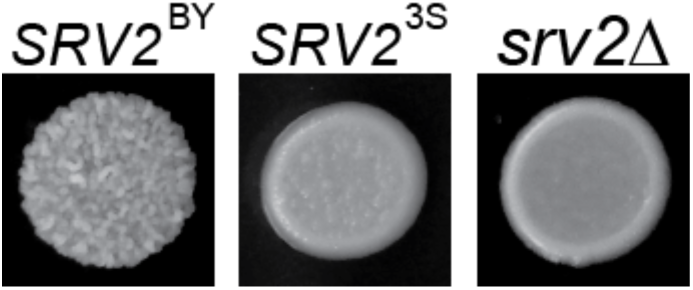
The *SRV2* gene underlies a locus on chromosome XIV that interacts with GPb. Replacement of the BY *SRV2* allele with the 3S version results in loss of the rough phenotype. A similar loss of phenotype is also observed when *SRV2* is deleted.

**Figure S7:**
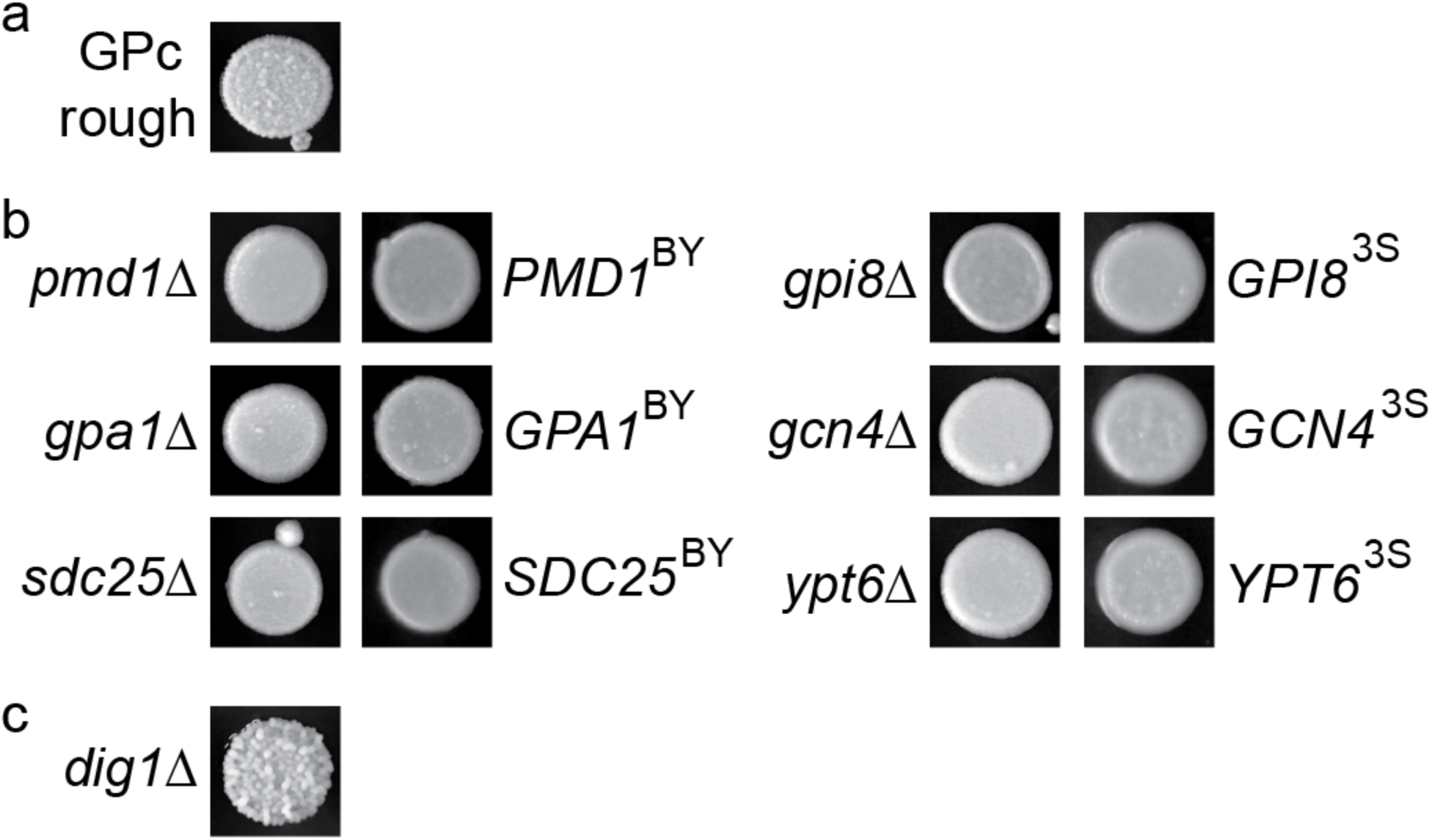
Genetic engineering supports interactions between GPc and identified candidate genes. Complete deletion of the candidate genes in a GPc rough segregant was performed using tailed *kanMX* cassettes. The phenotype of the GPc rough segregant is shown in **(a)**. In **(b)**, we show that deletion of six candidate genes causes loss of the phenotype. We also show that allele replacement with the non-causal allele at these genes results in no phenotypic recovery, implying these genes harbor functional genetic variation. Note, these replacements involved the entire coding region, as well as 100 to 200 bases of promoter sequence and 100 to 200 bases of downstream sequence. Control experiments were performed in parallel and resulted in restoration of the phenotype. In **(c)**, for the seventh candidate gene, deletion enhances the phenotype. Notably, *DIG1* encodes a repressor of the *FLO11* activator Ste12. Thus, the phenotypic enhancement seen upon its deletion is consistent with cryptic variation promoting transcription of *FLO11*.

**Note S1:** For GPb individuals expressing the rough phenotype at 21°C, we detected two multi-locus genotypes among segregants from the backcross to 3S. We also identified a third genotype in the second-generation backcross population, involving a different combination of alleles than those from the initial 3S backcross.

**Note S2:** We deleted each non-essential gene at the GPb chromosome II locus – *RRT2*, *HIS7*, *ARO4*, *MRPS5*, *MTC4*, *SHG1*, and *YBR259W* – and found that none of these deletions caused loss of the rough phenotype. This implies that loss-of-function in a non-essential gene is not responsible for the trait at this locus, and the causal genetic variant is either a gain-of-function polymorphism or is an essential gene. Among the essential genes at this locus, *SRB6* was determined to be the strongest candidate.

